# Preclinical efficacy of bacosides-lauric acid nano-herbal formulation in comparison to rivastigmine for treatment of Alzheimer’s disease

**DOI:** 10.1101/2022.06.25.497622

**Authors:** Ashish Kumar, Kritika Goyal, Muneesh Pal, Prabhat Upadhyay, Sarika Gupta, Veena Koul, Arpita Konar

**Author notes:** Correspondence: Ashish Kumar,; Arpita Konar,; Veena Koul,; Sarika Gupta. These authors have contributed equally to this work (Kritika Goyal, Muneesh Pal).

## Abstract

Despite decades of rigorous scientific endeavors for Alzheimer’s disease (AD) drug development, massive failures in clinical trials is continuously posing healthcare and societal burden. Currently recommended single targeted drugs including rivastigmine with limited bioavailability can alleviate AD symptoms only for a limited period of time but unable to reverse the disease progression. Recent evidences consider poly-pharmacological and multi-targeted agents aiming at amyloid and tau burden, neuroinflammation, neuroprotection and cognitive enhancement as potential treatment options. In this regard, bioactive herbal compounds with holistic action and minimal adversities have gained prominence, though lacunae in their scientific validation and limited bio-permeability to cross BBB represent major hurdles. Previously, we showed that lactoferrin conjugated PEG-S-S-PLA-PCL-OH efficaciously delivered herbal compounds to brain and such nano-herbal formulation of bacosides-lauric acid (BAN-LAN) attenuated neuronal damages induced by scopolamine *in vitro*. Here, we tested the preclinical potential of BAN-LAN in reversal of AD pathologies in 5XFAD transgenic mice. Our nano-herbal formulation substantially reduced amyloid burden by clearing Aβ plaques in the hippocampus of 5XFAD mice and also attenuated aβ42 induced alterations in AD associated gene expression in hippocampal neurons *in vitro*. It showed neuroprotection by rescuing neuronal damage and promoting neurogenesis in the hippocampus of AD transgenic mice. Anti-neuroinflammatory properties were also exhibited by the formulation as evident from inhibition of hippocampal astrocytic and microglial activation in 5XFAD mice. BAN-LAN showed cognitive efficacy by restoring memory impairment in AD transgenic mice and effects were more pronounced than unconjugated natural form and rivastigmine. These findings suggest that BAN-LAN may serve as a promising therapeutic agent for AD with better brain penetration and targeting multiple AD pathways.

## Introduction

Alzheimer’s disease (AD), a progressive neurodegenerative pathological condition and most prevalent dementia presents a global epidemic with enormous healthcare, economic and societal burden. It is estimated that over 55 million worldwide population is affected with AD and other forms of dementia. Epidemiological studies have suggested that around 150 million people will suffer from AD by 2050 (Policarpo and d’Ydewalle, 2022). AD primarily results in memory, thinking and other cognitive deficits (Perkovic et al., 2018) that initiate slowly and deteriorates with time, severely interfering with tasks of daily life (Alzheimer’s disease facts and figures, 2022). It is characterized by neuronal death and synaptic loss compromising cognitive functions in brain regions primarily the hippocampus (Hadipour et al., 2021). Various complex mechanisms are involved in the progression of AD (Guo et al., 2020). Despite extensive information on AD pathophysiology, hallmarks, risk factors and expensive clinical trials, this alarming neurological ailment is still incurable.

Currently available limited therapeutic options include acetylcholinesterase inhibitors (rivastigmine, galantamine, tacrine, donepezil) for any stage of the disease and NMDA receptor antagonist memantine for moderate to severe clinical condition. However, these drugs only provide symptomatic relief and several patients do not respond to them owing to their single target action, low bioavailability across the blood brain barrier and individual heterogeneity in clinical profiles (Casey et al., 2010; Abeysinghe et al., 2020). Recently, the FDA approved Aducanumab is the first drug targeting the causative pathway of AD mainly amyloid β (Aβ), though therapeutic efficacy is not yet established and still there is a pressing need to develop novel AD therapies to reverse as well as retard the progression of disease (Rabinovici, 2021).

In the last decade, more than 200 investigational programs on AD therapeutics have either been abandoned during the developmental phase or failed in clinical trials. Inadequate understanding of the complex pathophysiology of AD and aiming only at reduction of Aβ pathological aggregates or phosphorylated tau has been one of the key reasons for such failures. Moreover toxic side effects, inappropriate dosage and timing of treatment, and reduced bioavailability have added to the obstacles (Yiannopoulou et al., 2019). Recent researches have revealed that apart from amyloid and tau burden, neuroinflammation, neuronal atrophy, loss of synaptic connectivity play crucial role in the pathogenesis and progression of AD (Xie et al., 2022). Therefore, multi-targeted and combinatorial therapies capable of reversing AD pathology as well as maintaining body homeostasis are considered as prospective treatment approach.

Neuroactive herbal compounds holds great promise in AD and related neurodegenerative diseases due to their holistic action, less toxicity with preventative as well as curative abilities (Wadhwa et al., 2016). However, herbal compounds have high molecular weight with complex chemical composition and less solubility in water making their bioavailability difficult and challenging. Nanoparticle-guided drug delivery of these compounds can enhance their bioavailability and prove effective in AD therapeutics. Earlier, we showed that lactoferrin conjugated PEG-S-S-PLA-PCL-OH polymersomes efficaciously delivered herbal compounds, bacosides (BAN) to brain and it alleviated scopolamine induced memory impairment better than its unconjugated natural form (BA) (Goyal et al., 2018). Considering greater therapeutic potential of combinatorial herbal formulations, we also investigated the co-administering effect of BAN and nano-formulation of coconut oil derived lauric acid (LAN) on scopolamine challenged neurons. Nano-herbal formulation of BAN-LAN in 1:1 ratio, significantly attenuated scopolamine induced neuronal damages *in vitro*.

Here, we evaluated the potential of BAN-LAN in reversal of AD pathologies using 5XFAD transgenic mice. We also compared the therapeutic efficacy of BAN-LAN with FDA approved drug rivastigmine. BAN-LAN substantially cleared amyloid plaques, rescued neuronal damage and induced neurogenesis in the hippocampus of 5XFAD mice. AD associated gene expression changes were also attenuated upon treatment with BAN-LAN formulation. BAN-LAN treatment inhibited astrocytic and microglial activation, characteristic of neurodegenerative insult thereby indicating its anti-inflammatory property. Neuroprotective effect of BAN-LAN was accompanied by its ability to ameliorate cognitive impairment as it significantly recovered spatial memory loss of 5XFAD mice. Effects of BAN-LAN were more pronounced as compared to their unconjugated natural form and nano-encapsulated rivastigmine (RiN) treated groups in all the cellular, molecular and behavioral parameters assessed. These findings suggest that BAN-LAN may serve as a promising therapeutic agent for AD with better brain penetration and targeting multiple AD pathways. However, translational potential of this work necessitates investigation in human models of AD.

## Materials and methods

### Synthesis of PEG-S-S-PLA-PCL-OH: the triblock copolymer

The detailed synthesis procedure has been explained in our previous research article (Goyal et al., 2018). Briefly, the novel triblock copolymer was developed in four steps (Fig.1). In first step, terminal hydroxy (-OH) group of PEG was reacted with succinic anhydride to synthesize PEG-COOH. Disulfide linkage was introduced in PEG-COOH by using 2-hydroxyethyl disulfide in the following step. PEG-COOH with disulfide linkages i.e. PEG-S-S-OH was employed as initiator for the ROP (ring opening polymerization) of lactide. Further, PEG-S-S-PLA-OH was used as an initiator for the ROP of ε-caprolactone and finally PEG-S-S-PLA-PCL-OH was formed (Kumar et al., 2015).

**Fig. 1.**
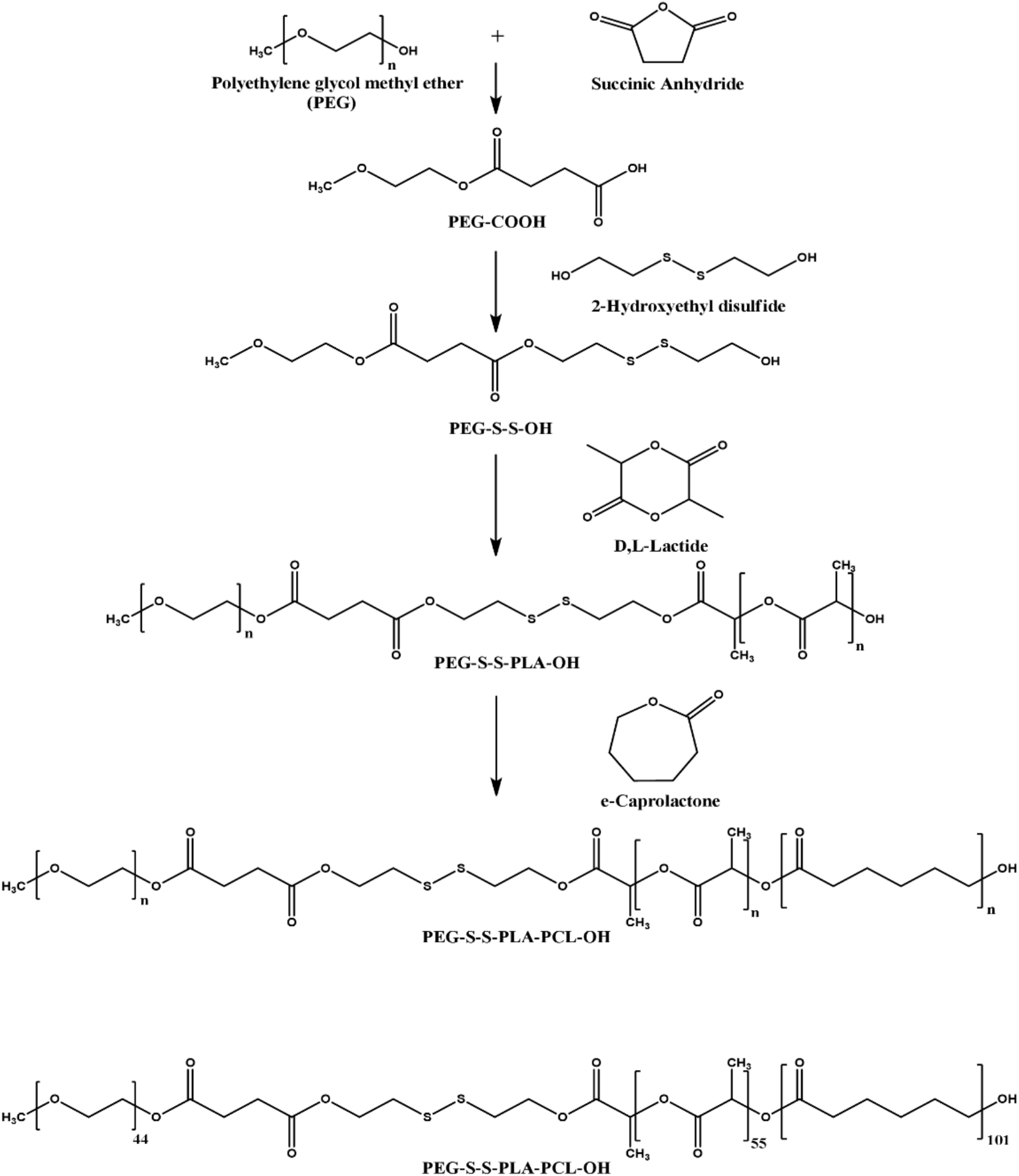
PEG-PLA-PCL-OH triblock copolymer synthesis by ring opening polymerization. [Reproduced from Goyal et al., (2018) with permission from the Royal Society of Chemistry].

### Conjugation of Lactoferrin (Lf) to triblock copolymers

Lf (2 mg) was activated in DMSO (10 mL) for 100mg polymer (0.0059 moles) by EDC (2equivalents of the polymer, 2.28 mg) and NHS (2 equivalents of the polymer, 1.35 mg) for 12 hours under nitrogenous condition. The polymersomes were added to the system and the reaction was further allowed to proceed for 12 hours under similar conditions. On completion of the reaction unreacted Lf, NHS and EDC were eliminated by dialysis. The polymersomes were characterized by assessing the change in zeta potential, nitrogen content and circular dichroism.

### Preparation of bacosides encapsulated nanoparticles (BAN), Lauric Acid encapsulated nanoparticles (LAN) and rivastigmine encapsulated nanoparticles (RiN)

Briefly the active components of *B. monniera* were separated from the crude extract by butanolic method (Sharath et al., 2010). The butanolic fraction was separated out and precipitated in ethyl acetate twice. The active components were recovered in methanol, this was termed as the bacoside fraction, which will be addressed as ‘bacosides’. The detailed methodology has been published in our pervious publication (Goyal et al., 2018). Polymersomes loaded with Bacosides were prepared by an altered version of the o/w emulsion solvent evaporation process. The triblock copolymer and Bacosides (120mg/kg body weight) (Bacoside: polymer ratio 1:2) were dissolved in methanol-dichloromethane mixture (1:2) which made the organic phase. The organic phase was emulsified with aqueous phase containing Pluronic F-68 2% using an ultra-probe sonicator (UP200S Hielscher) at an output of 40 W for 5 min in an ice bath. The organic phase was evaporated by keeping it overnight on a magnetic stirrer. The resulting Bacoside loaded polymersomes suspension was ultra-centrifuged (3K3O, Sigma) at 13,000xg rpm for 30 min at 4°C (123, 124).

Similarly polymeric nanoparticles encapsulated with lauric acid (LA) were prepared by solvent-diffusion method. LA and the triblock co-polymer were co-dissolved in DMSO in the ratio 1:10. The solution was slowly added to deionized water containing 0.1% Pluronic F62 as a stabilizing agent. The mixture was homogenized well and the resulting nanoparticles were dialyzed three times against PBS (pH 7.4) for 2 h and then stored at 4°C for further use. Size distribution and zeta potential were measured by Zeta Sizer. RiNs were prepared by nanoprecipitation method where rivastigmine and the tri block co-polymer were dissolved in acetone, this was slowly added to aqueous phase supplemented with Pluronic F62. The organic solvent was then allowed to evaporate for 4 h with continuous stirring on a magnetic stirrer. The resulting suspension was centrifuged at 25,000 rpm for 1 h at 4 °C using high-speed centrifuge (Sigma) (Wilson et al., 2008).

### Preparation of Amyloid beta 1-42 oligomers

Preparation of oligomeric amyloid β (aβ)1-42 was done from lyophilised aβ 1-42 powder(Plz mention source) as mentioned by Fa et al. (2010). Briefly, the lyophilised powder was resuspended in ice cold 1,1,1,3,3,3-Hexfluoro-2-propanol (HFIP) and incubated for 2 h at room temperature to allow the aβ 1-42 monomerisation. Further, the aβ 1-42 /HFIP solution containing vials were incubated in speedVac centrifuge under vacuum to obtain the peptide film. This film was resuspended in DMSO and 5 mM aβ 1-42 solutions were made. In order to oligomerise the aβ 1-42 solution, it was diluted with sterile Phosphate buffer and incubated at 4°C for 12 h.

### Animal

Primary neuronal culture studies were performed using Swiss albino mice maintained in 12 h light/ dark cycle and free access to food and water *ad-libitum* in the Central Animal House of All India Institute of Medical Sciences (AIIMS), New Delhi, India. Experiments were conducted as per the guidelines and approval of the institutional animal ethical committee (ethical approval number 119/ IAEC-1/ 2019). *In-vivo* studies were performed using 5XFAD mouse model after approval from the Institutional Animal Ethics Committee IAEC#565/20 of National Institute of Immunology and according to ARRIVE (Animal Research: Reporting of In-Vivo Experiments) guidelines. The transgenic mice were procured from Jackson’s laboratory and maintained at the animal facility of National Institute of Immunology, New Delhi.

### Genotyping

The mice were segregated into wild type and 5XFAD transgenic using genotyping. The DNA was isolated from the tail of these animals using DNA extraction kit. APP and PSEN genes were screened in these animals using specific primers (Table-1). Mice expressing both the genes were considered as transgenic and others as wild type. The protocol for DNA extraction was followed as per the manufacturer’s instructions.

**Table-1.**
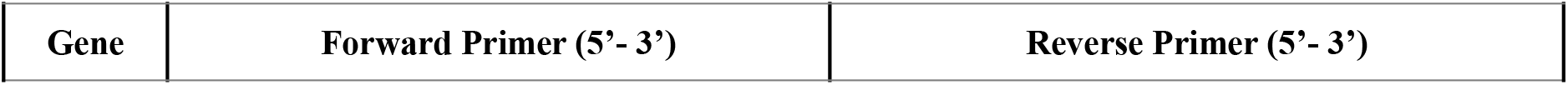
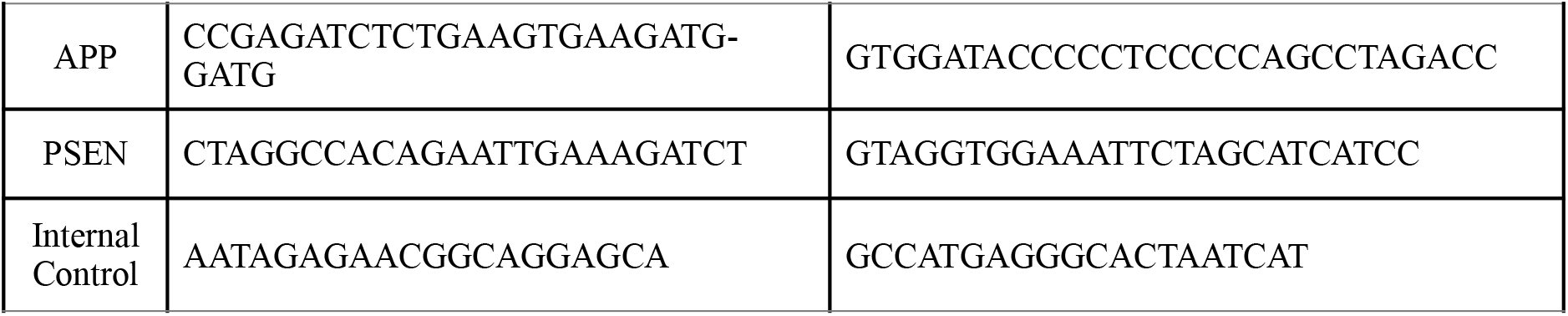
List of primer sequences used for genotyping of 5XFAD transgenic mice model.

### Cell culture

#### Primary culture of mouse hippocampal neurons

Primary cultures of hippocampal neurons were prepared from embryonic day (E) 18 mice. Hippocampi were dissected out and subjected to mechanical dissociation followed by trypsin digestion [0.02% (w/v)] at 37 °C for 10 min and inactivated by FBS. The dissociated cells were triturated and single cell suspension was prepared. Cells were seeded in complete neurobasal medium supplemented with 2% B27 and 2mM glutamax and incubated in a humidified 5% CO2/95% air atmosphere at 37 °C (Cardenas-Aguayo Mdel et al., 2013).

### Drugs and treatments

Primary hippocampal neurons (DIV5), 60–70% confluent were subjected to various drug treatments. The treatment groups included drug vehicles (0.9% saline; 0.05% Tween 80), nanoparticle alone (N), aβ 1-42 oligomers, aβ 1-42 oligomer co-administered with 8 different formulations comprising of Ri, RiN, BA, BAN, LA, LAN, BA+LA (1:1) and BAN+LAN (1:1). Cells were treated for 48 h and subjected to cellular and molecular assays. 5XFAD mice were treated with drug vehicles (0.9% saline; 0.05% Tween 80), N, RiN, BA+LA (1:1) and BAN+LAN (1:1). Wild type control was additionally taken. The animals were treated for 45 days, and thereafter, Morris water maze test was conducted to evaluate learning and memory of the mice.

### Morris water maze

The Morris water maze (MWM) was used to assess spatial memory in AD transgenic mice with respect to the wild type mice. In MWM paradigm, three trials of 60 s with 15 min interval between trials were given for five consecutive days followed by probe trial. The stage position was kept different and the trial was declared completed when the mice reached the platform or the time was over. In case the mice could not find the platform, it was directed towards the platform and trained to sit on the platform for 10s in the training phase. For probe trial, the platform was removed and the mice was placed in opposite quadrant to the quadrant where the platform was kept in acquisition testing. The water in MWM pool was kept opaque and ANY-MAZE software was used for data acquisition and analysis.

### Nissl and Congo red staining

Nissl staining was performed to assess neuronal loss and recovery by drug formulations. Briefly, mouse brain frozen coronal sections were hydrated in graded ethanol solutions (100–70%) to distilled water for 30 seconds each. Thereafter, sections were stained with 1% cresyl violet in 70% ethanol for 1 min, immersed in distilled water, dehydrated with increasing concentrations of ethanol (70–100%), cleared in xylene, counter stained with hematoxylin and mounted on coverslips. Images were captured using bright field microscope and Nissl positive cells were counted using ImageJ software.

Congo red staining was also performed using coronal sections of 5XFAD and WT mouse brain. Briefly, the sections were washed with 1XPBS and permeabilized with 0.1% Triton-X100 in PBS. The sections were then incubated in Congo red solution for 60 min and differentiated in alkaline alcohol solution (1% Sodium hydroxide, 50% alcohol). The sections were dipped in ammonia water for 30 sec and dehydrated through graded alcohol series. Further, the sections were cleared in xylene, mounted with resinous mounting medium and images were captured.

### Immunofluorescence

Immunofluorescence analysis was performed using hippocampal sections from 5XFAD and WT mice. Briefly, the cryosections were washed with warm 1XPBS and permeabilized with 0.1% Triton-X100 in PBS (PBST) for 45 mins. Thereafter tissue sections were blocked for 2h at room temperature followed by incubation overnight with primary antibodies {1: 500 dilution of Anti-GFAP antibody (ab7260), Abcam and 1:200 dilution of Rabbit anti Iba1(019-19741) antibody, Wako}. After incubation, the sections were washed three times with 0.1% Tween 20 in PBS, for 10 min each. The sections were incubated with secondary antibodies {Alexa fluor 594 donkey anti-rabbit IgG (A21207) for GFAP and Alexa fluor 488 donkey anti-rabbit IgG (A21206) for Iba1}for 1 h at room temperature, washed three times, counterstained with DAPI and photomicrographs were captured by FLoid fluorescence microscope. To quantify the immunoreactivity of GFAP and Iba1, corrected total cell fluorescence {CTCF= Integrated Density – (Area of selected cell X Mean fluorescence of background readings)} was measured using ImageJ software.

### Quantitative real-time PCR (qRT-PCR)

Total RNA was isolated from control and experimental groups of mouse primary hippocampal neurons and 2 μg of RNA from each group was reverse transcribed to cDNA using cDNA Synthesis Kit-AB1453B (Thermo). RT-PCR was carried out using SYBR Green master mix (Sybr Green-A6001 Promega) for detection in Light cycler LC 480 (Roche). All primers used for qRT-PCR are given in Table-2. The endogenous control 18S was used to normalize quantification of the mRNA target. In order to analyze RT-PCR data, the 2^-ΔΔCt value was used to calculate relative fold change in mRNA expression and plotted as histograms.

**Table-2.**
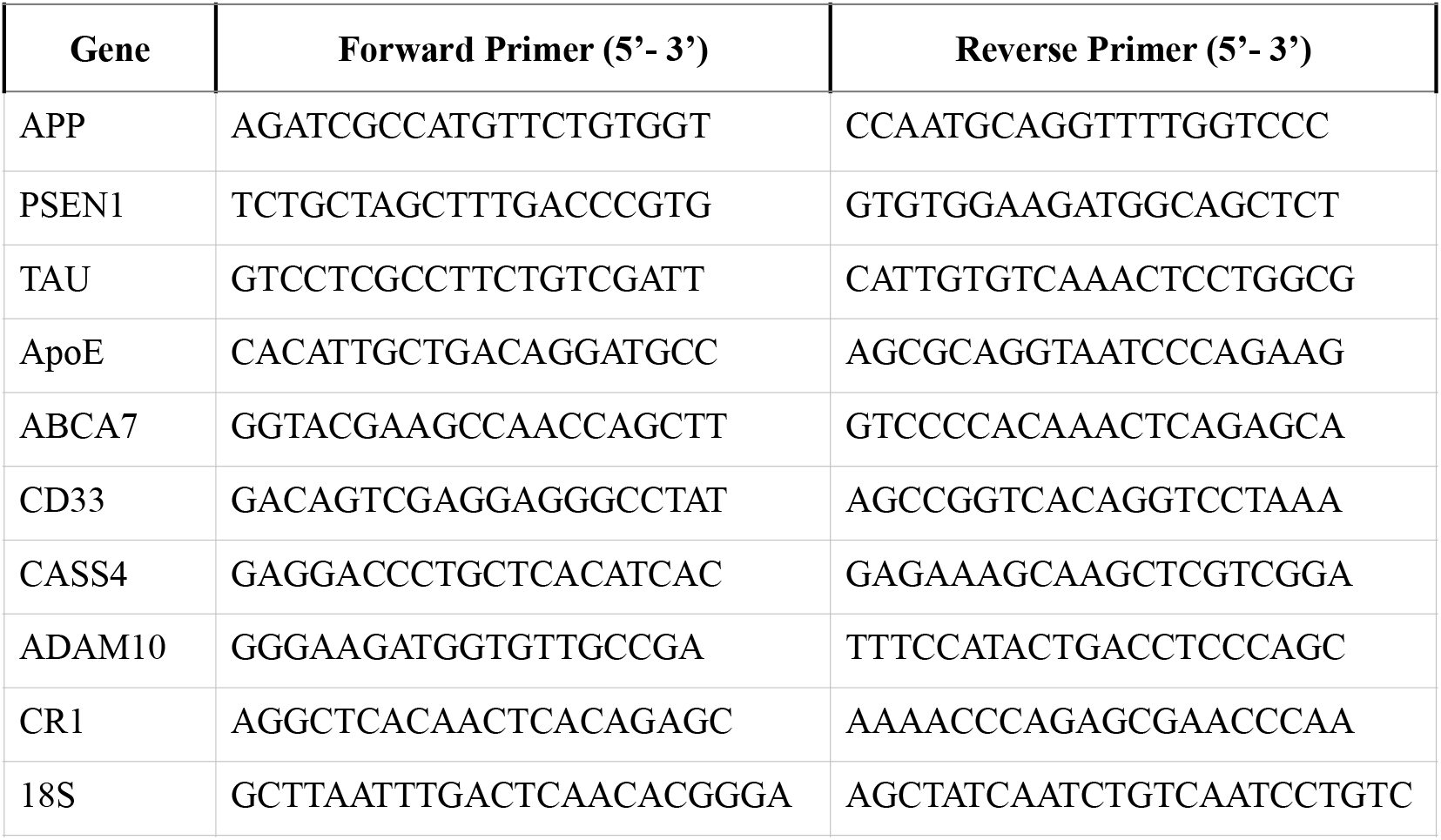
List of primer sequences used for qRT-PCR analysis.

### Statistical analyses

All statistical analyses were performed using Prism 8 (GraphPad Software). Sample sizes, statistical analysis used and exact p values for each experiment is mentioned in respective figure legend section. In all the *in vitro* studies, treatments were performed in three independent culture dishes, experiment was repeated three times and images were captured from random fields. For multiple comparisons, one way analysis of variance (ANOVA) and Brown-Forsythe tests were run (p< 0.05) between the groups. Total number of animals used in experiment and number of animals in each experimental group is also mentioned. Each animal was considered as a biological replicate and all our replicates presented in figures were biological. Experiment repeated with same samples was considered as technical replicate. In case of Nissl and Congo red staining, immunofluorescence and qRT-PCR experiments three biological replicates were further run in three technical replicates. For Morris water maze test, the experiment was repeated three times with n=6 mice/group. Histograms were represented as mean of the data (±SD) and statistical significance was calculated using One way ANOVA followed by Brown-Forsythe test (p< 0.05) between the groups.

## Results

### BAN-LAN attenuated AD associated gene expression changes in aβ42 induced hippocampal neurons *in vitro*

BAN-LAN formulation used in these experiments has been characterized for its physico-chemical and biological properties including brain targeted delivery in our previous reports (Goyal et al 2018, 2020, 2021). Here, we evaluated the effect of BAN-LAN treatment on the expression of genes associated with AD in primary hippocampal neuronal culture induced by aβ42. The analysis showed that Aβ42 upregulated all the genes (ABCA7 2.35-fold; ADAM10 2.85-fold; ApoE 2.85-fold; BAK 3.81-fold; CASS4 2.57-fold; CD33 3-fold; CR1 1.79-fold; MAPT 3.44-fold; PS1 2.26-fold) but downregulated APP (0.3-fold) in primary neurons. BAK and MAPT gene expression were most induced by Aβ42. Treatment with both the herbal compounds BA (ABCA7 0.86-fold; ADAM10 1.61-fold; ApoE 1.61-fold; APP 0.73-fold; BAK 2.11-fold; CASS4 0.92-fold; CD33 1.41-fold; CR1 1.13-fold; MAPT 1.19-fold; PS1 0.59-fold) and LA (ABCA7 1.06-fold; ADAM10 0.22-fold; ApoE 0.22-fold; APP 0.71-fold; BAK 2.5-fold; CASS4 1.14-fold; CD33 1.4-fold; CR1 1.01-fold; MAPT 1.15-fold; PS1 0.63-fold) alleviated the changes. However, their respective nano-conjugates of BAN (ABCA7 0.74-fold; ADAM10 0.50-fold; ApoE 0.50-fold; APP 0.95-fold; BAK 0.70-fold; CASS4 0.64-fold; CD33 0.77-fold; CR1 0.98-fold; MAPT 0.79-fold; PS1 0.26-fold and LAN (ABCA7 0.76-fold; ADAM10 0.18-fold; ApoE 0.18-fold; APP 0.92-fold; BAK 0.74-fold; CASS4 0.68-fold; CD33 0.81-fold; CR1 0.81-fold; MAPT 0.81-fold; PS1 0.28-fold) were more effective in attenuating the effects of Aβ42. BA-LA cocktail in equal ratio (ABCA7 0.46-fold; ADAM10 0.42-fold; ApoE 0.42-fold; APP 1.26-fold; BAK 2.04-fold; CASS4 0.51-fold; CD33 0.49-fold; CR1 0.93-fold; MAPT 0.48-fold; PS1 0.50-fold) was even more potent than individual compounds. Nano-formulation of pharmacological compound RiN (ABCA7 1.25-fold; ADAM10 1.08-fold; ApoE 1.07-fold; APP 0.63-fold; BAK 1.14-fold; CASS4 1.42-fold; CD33 1.69-fold; CR1 1.42-fold; MAPT 1.48-fold; PS1 1.04-fold) also altered gene expression but to a lesser extent than herbal compounds for majority of the genes tested. BAN-LAN cocktail showed the highest promising effects amongst all drug formulations (ABCA7 0.22-fold; ADAM10 0.13-fold; ApoE 0.13-fold;

### BAN-LAN diminished Aβ burden efficaciously than RiN in the hippocampus of 5XFAD mice

Aβ deposition and formation of senile plaques is a major hallmark of AD. In order to assess whether BAN-LAN treatment affects Aβ deposition in AD mice, we performed Congo red staining for visualizing the amyloid plaques. Hippocampal region of 5XFAD mice (Tg) showed significantly higher levels of aβ senile plaques as compared to wild type (WT) control counterparts. (Fig. 3). As anticipated, nano formulation of approved drug rivastigmine (RiN) lowered amyloid plaque burden in Tg mice as compared to WT control mice. However, BA-LA combination as well as its nano-formulation, BAN-LAN was more efficacious in clearing the aβ plaques. More importantly BAN-LAN showed the most promising effects amongst all the drug treated groups. (Fig.3).

**Fig. 2.**
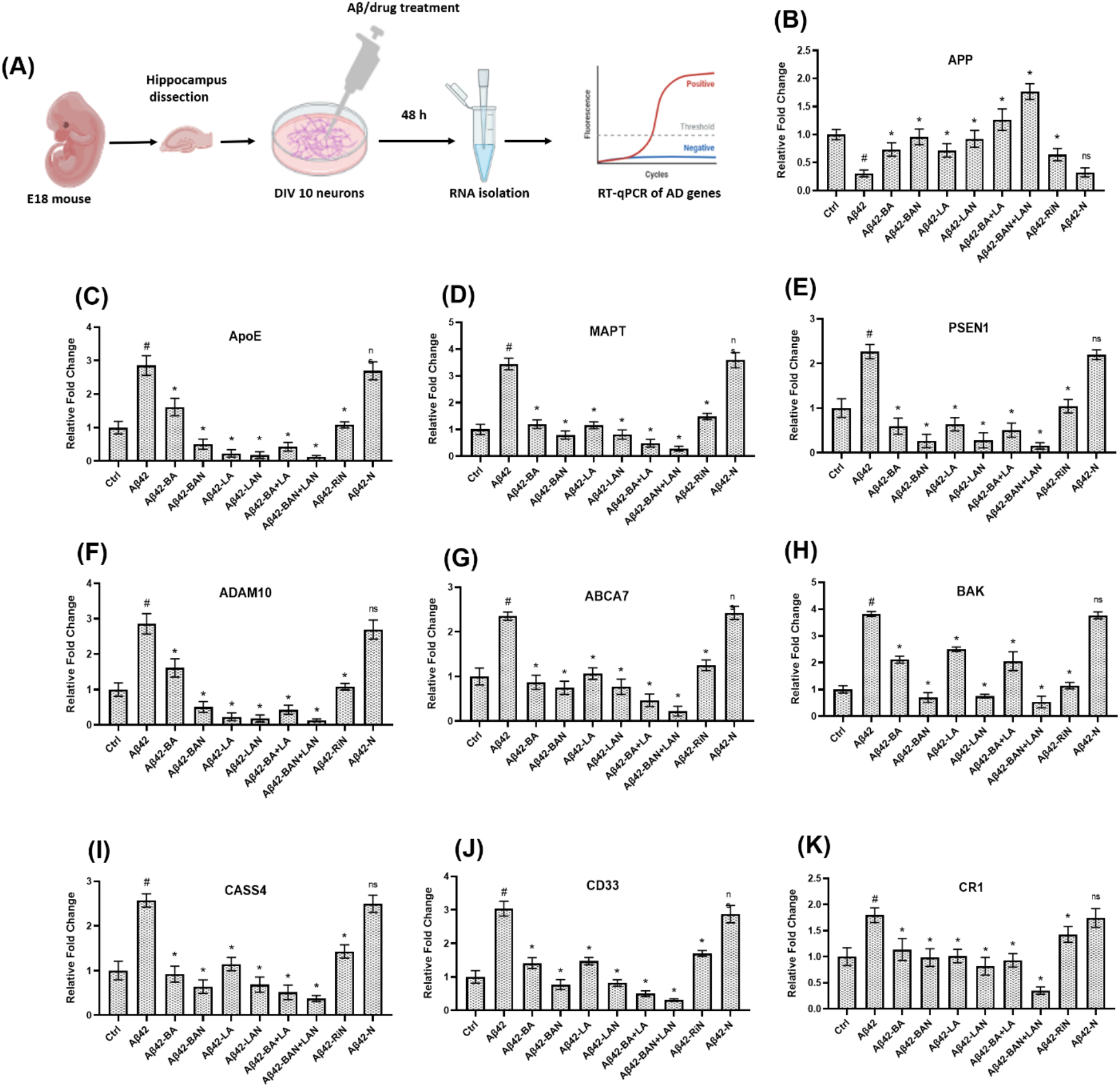
Attenuation of AD associated gene expression changes by nano-herbal formulation of bacosides and lauric acid better than rivastigmine in aβ42 induced mouse primary hippocampal neurons. (A) Experimental timeline of mouse primary hippocampal neurons culture. Expression analysis of AD associated genes (B) APP, (C) ApoE, (D) MAPT, (E) PSEN1, (F) ADAM10, (G) ABCA7, (H) BAK, (I) CASS4, (J) CD33, and (K) CR1.The treatment groups includes primary hippocampal neurons treated with *group-1* drug vehicles (Control), *group-2* aβ 1-42 oligomers (Aβ42), *group-3* aβ 1-42 oligomer co-administered with bacosides (Aβ42-BA), *group-4* aβ 1-42 oligomer co-administered with nano-herbal formulation of BA (Aβ42-BAN), *group-5* aβ 1-42 oligomer co-administered with lauric acid (Aβ42-LA), *group-6* aβ 1-42 oligomer co-administered with nano-herbal formulation of LA (Aβ42-LAN), *group-7* aβ 1-42 oligomer co-administered with BA and LA (Aβ42-BA+LA), *group-8* aβ 1-42 oligomer co-administered with BAN and LAN (Aβ42-BAN+LAN), *group-9* aβ 1-42 oligomer co-administered with nano-formulation of rivastigmine (Aβ42-RiN) and *group-9* aβ 1-42 oligomer co-administered with nanoparticle alone (Aβ42-N). Histogram represents mean of the data (±SD). Statistical analysis were performed using One way Anova followed by Brown-Forsythe test (p< 0.05) between the groups (# -Aβ42 vs Control, * - other treatment group vs Aβ42 and ns – non significant). APP 1.76-fold; BAK 0.53-fold; CASS4 0.37-fold; CD33 0.30-fold; CR1 0.35-fold; MAPT 0.28-fold; PS1 0.15-fold) prompting us to study it in depth in AD *in vivo* model system (Fig.2).

**Fig. 3.**
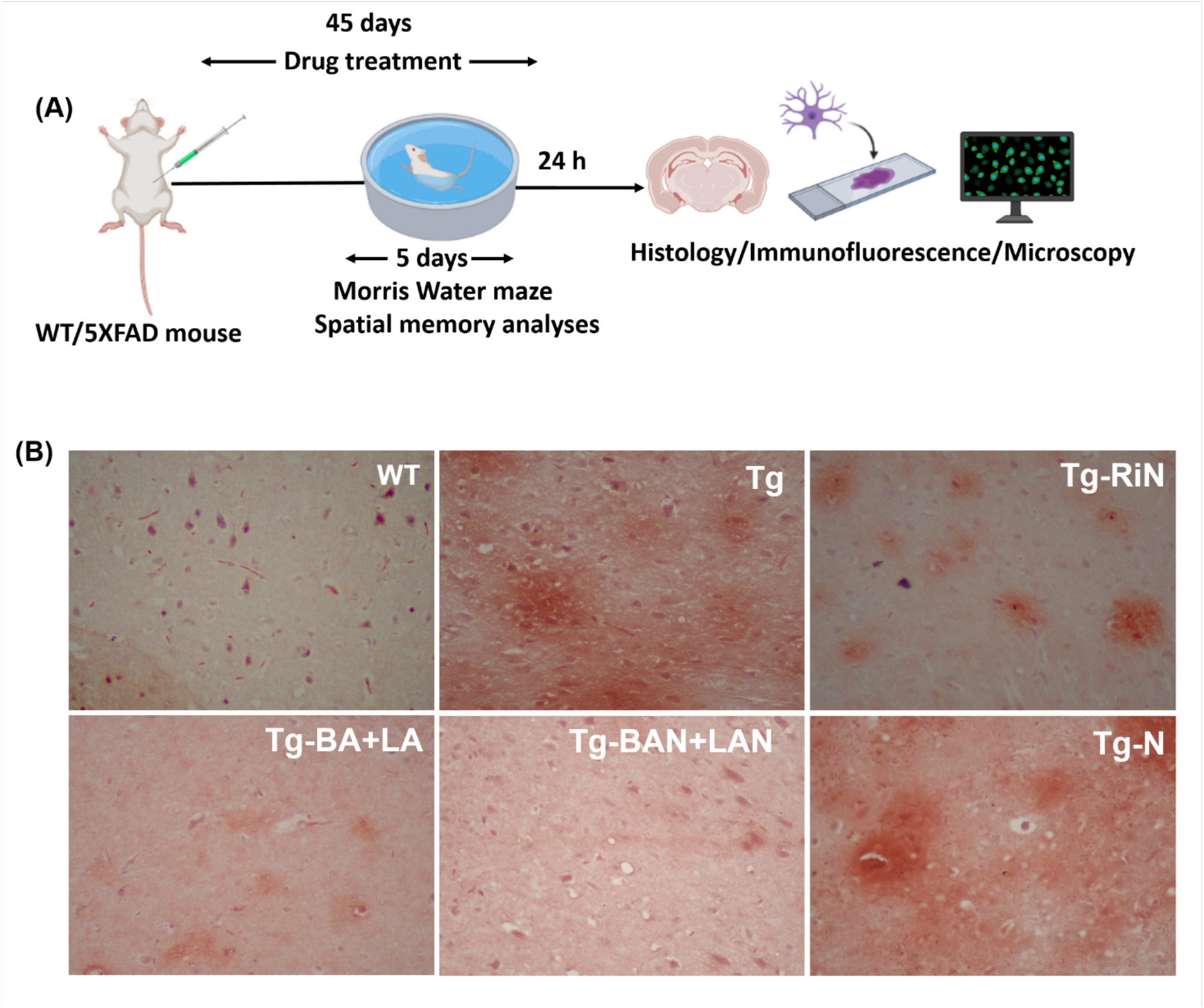
Reduction in aβ 1-42 burden by nano-herbal formulation of bacosides and lauric acid efficaciously than nano-herbal formulation of rivastigmine in the hippocampus of 5XFAD. (A) Experimental timeline of drug treatment in 5XFAD mice. (B) The treatment groups includes wild type (WT) and 5XFAD mice. G*roup-1* Wild type treated with saline (WT), *group-2*5XFAD mice treated with Saline (Tg), *group-3*Tg treated with nano-formulation of rivastigmine (Tg-RiN), *group-4* bacosides and lauric acid (Tg-BA+LA), *group-5*Tg treated with nano-herbal formulation of BA and LA (Tg-BAN+LAN) and *group-6* Tg treated with nanoparticle alone (Tg-N). Histological analysis of aβ 1-42 deposits in the coronal sections from hippocampal region. The sections were stained by Congo red and counter stained by hematoxylin. (if possible include birefringes of congored to show amyloid burden).

### BAN-LAN reversed AD associated hippocampal neurodegeneration potently than RiN

Nissl staining was performed to investigate the extent of neurodegeneration in the hippocampus of 5XFAD transgenic (Tg) mice and to assess recovery potential of BAN-LAN and other drug formulations. Tg mice showed obvious neuronal damage throughout the hippocampus evident from reduced number of Nissl positive neurons showing 65% loss in CA1, 67% in CA3 and 53% in DG as compared to 100% in WT. In CA1 and CA3 sub-regions, RiN and BA-LA treatment significantly reversed the loss by maintaining neuron count similar to WT levels while BAN-LAN even increased the number of Nissl positive neurons more than WT (CA1-123% and CA3-150%). All the drug formulations increased neuron number (RiN 120%; BA-LA 130%; BAN-LAN 144%) as compared to WT in DG. Overall, BAN-LAN was most efficient in alleviating the neuronal damage and increasing the number of neuronal cells in the CA1, CA3 and DG regions (Fig.4).

**Fig. 4.**
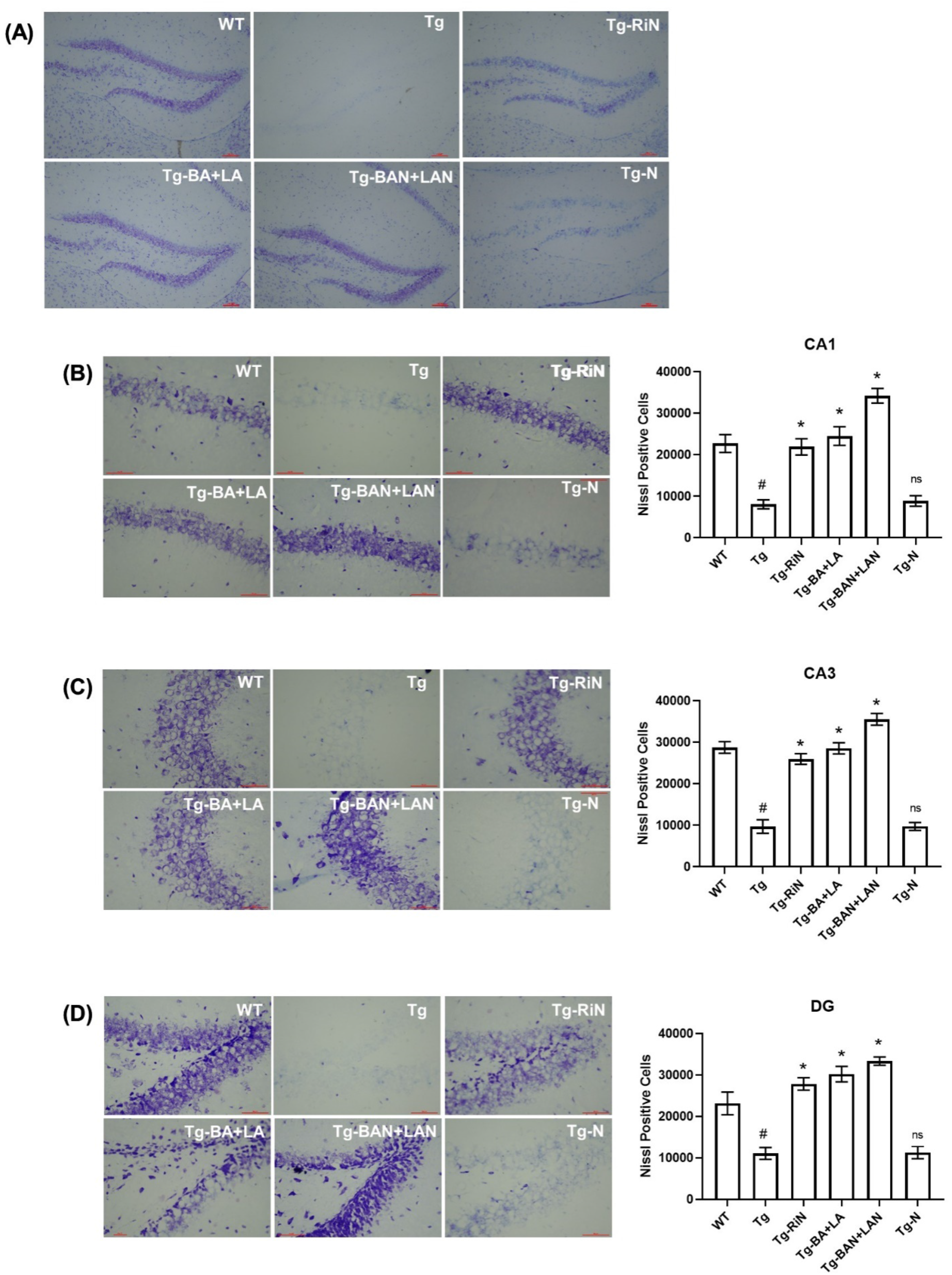
Reversal of AD associated hippocampal neurodegeneration by nano-herbal formulation of bacosides and lauric acid potently than nano-herbal formulation of rivastigmine in 5XFAD mice. Histological analysis of Nissl positive cells to understand neurodegeneration in the coronal sections from hippocampal region. Images represent (A) total hippocampus, (B) CA1 region, (C) CA3 region and (D) dentate gyrus stained with Nissl staining method. The treatment groups includes wild type (WT) and 5XFAD mice. G*roup-1* Wild type treated with saline (WT), *group-2*5XFAD mice treated with Saline (Tg), *group-3*Tg treated with nano-formulation of rivastigmine (Tg-RiN), *group-4* bacosides and lauric acid (Tg-BA+LA), *group-5*Tg treated with nano-herbal formulation of BA and LA (Tg-BAN+LAN) and *group-6* Tg treated with nanoparticle alone (Tg-N). Histogram represents mean of the data (±SD). Statistical analysis were performed using One way Anova followed by Brown-Forsythe test (p< 0.05) between the groups (# - Tg vs WT, * - other treatment group vs Tg).

### BAN-LAN suppressed neuroinflammation by counteracting reactive astrogliosis and microglial over activation

Astrocyte activation or astrogliosis is one of the characteristic features of AD in response to accumulation of aβ plaques. We found dramatic astrocyte activation evident from increased GFAP immunofluorescence intensity in hippocampal CA1 (2.05-fold), CA3 (1.32-fold) and DG (2.43-fold-fold) region of 5XFAD transgenic (Tg) mice as compared to their age-matched wildtype (WT) littermates. BAN-LAN treatment (Tg-BAN+LAN) greatly controlled such astrocytic activation by decreasing GFAP levels with subtle regional differences being 0.51-fold in CA1, 0.15-fold in CA3 and 0.63-fold in DG. BA-LA (Tg-BA+LA) also dampened the astrocytic activation in all three sub regions of CA1 (0.78-fold), CA3 (0.23-fold) and DG (0.75-fold) but with lesser efficiency than BAN-LAN. Nano-rivastigmine (Tg-RiN) also counteracted astrogliosis but again with less potency than BAN-LAN in CA1 (1.02-fold) and CA3 (0.30-fold) and DG (0.89-fold) (Fig.5).

**Fig. 5.**
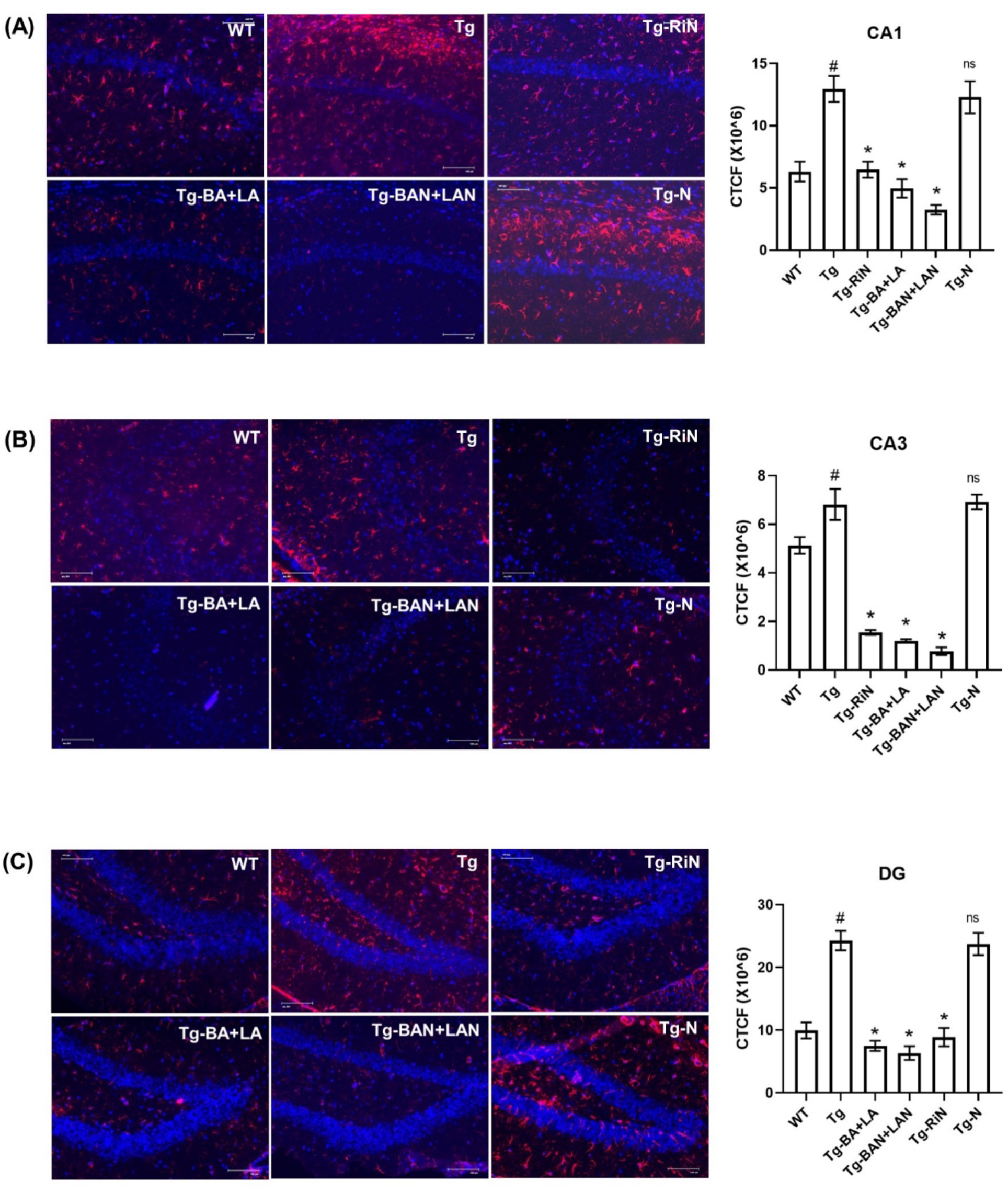
Suppression of reactive glia cells mediated neuroinflammation by nano-herbal formulation of bacosides and lauric acid efficaciously than nano-herbal formulation of rivastigmine in 5XFAD mice. Immunohistochemical analysis of glia positive cells (red) (mention 1° & 2° Antibody Company and dilution) count stained with nuclear stain DAPI to understand neuroinflammation in the coronal sections from hippocampal region. Images represent (A) CA1 region, (B) CA3 region and (C) dentate gyrus. The treatment groups includes wild type (WT) and 5XFAD mice. G*roup-1* Wild type treated with saline (WT), *group-2*5XFAD mice treated with Saline (Tg), *group-3*Tg treated with nano-formulation of rivastigmine (Tg-RiN), *group-4* bacosides and lauric acid (Tg-BA+LA), *group-5*Tg treated with nano-herbal formulation of BA and LA (Tg-BAN+LAN) and *group-6* Tg treated with nanoparticle alone (Tg-N).Histogram represents mean of the data (±SD). Statistical analysis were performed using One way Anova followed by Brown-Forsythe test (p< 0.05) between the groups (# - Tg vs WT, * - other treatment group vs Tg).

Iba-1 immunofluorescence staining was performed to evaluate aβ accumulation induced microglial over-activation in the hippocampus of Tg mice. Effect of BAN-LAN treatment along with comparative profiling of other drug formulations was also analyzed. We observed hyper activation of microglia determined by increase in Iba1 immunofluorescence intensity in the hippocampus (CA1 2.43-fold; CA3 2.28-fold; DG 2.19-fold) of Tg mice as compared to wildtype (WT) counterparts. All three drug formulations of RiN, BA-LA and BAN-LAN significantly relieved microglial overactivation but BAN-LAN showed highest efficacy particularly for CA3 (RiN 0.52-fold; BA-LA 0.56-fold; BAN-LAN 0.35-fold) and DG (RiN 0.69-fold; BA-LA 0.62-fold; BAN-LAN 0.28-fold) sub-regions (Fig.6).

**Fig 6.**
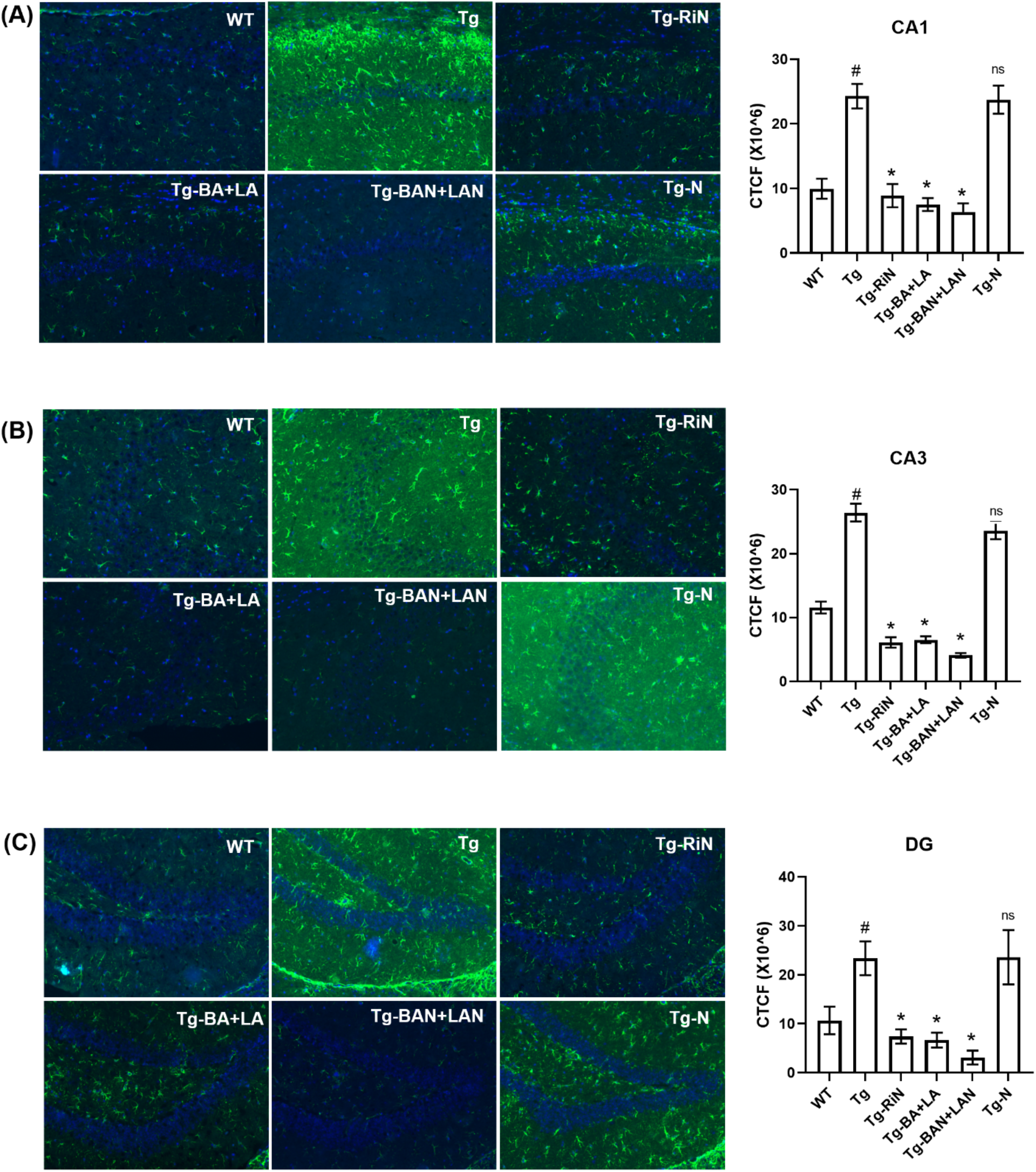
Suppression of microglial over activation mediated neuroinflammation by nano-herbal formulation of bacosides and lauric acid efficaciously than nano-herbal formulation of rivastigmine in 5XFAD mice. Immunohistochemical analysis of microglia positive cells (green) (mention 1° & 2° antibody company and dilution) count stained with nuclear stain DAPI to understand neuroinflammation in the coronal sections from hippocampal region. Images represent (A) CA1 region, (B) CA3 region and (C) dentate gyrus. The treatment groups include wild type (WT) and 5XFAD mice. G*roup-1* Wild type treated with saline (WT), *group-2*5XFAD mice treated with Saline (Tg), *group-3*Tg treated with nano-formulation of rivastigmine (Tg-RiN), *group-4* bacosides and lauric acid (Tg-BA+LA), *group-5*Tg treated with nano-herbal formulation of BA and LA (Tg-BAN+LAN) and *group-6* Tg treated with nanoparticle alone (Tg-N). Histogram represents mean of the data (±SD). Statistical analysis were performed using One way Anova followed by Brown-Forsythe test (p< 0.05) between the groups (# - Tg vs WT, * - other treatment group vs Tg).

### BAN-LAN ameliorated spatial memory impairment in 5XFAD mice

The effect of different treatment groups on the memory impairment of the AD transgenic mice was evaluated by Morris water maze. After training for 5 days, the AD transgenic mice showed a poor performance on finding the platform as compared to the wild type mice. Treatment of BA-LA, BAN-LAN or RiN significantly improved the memory impairment but the nano-herb formulations of BAN-LAN was more prominent in comparison to all the other treatment groups. It suggests that BAN-LAN has potential to ameliorate spatial learning and memory of the AD transgenic mice (Fig. 7).

**Fig 7.**
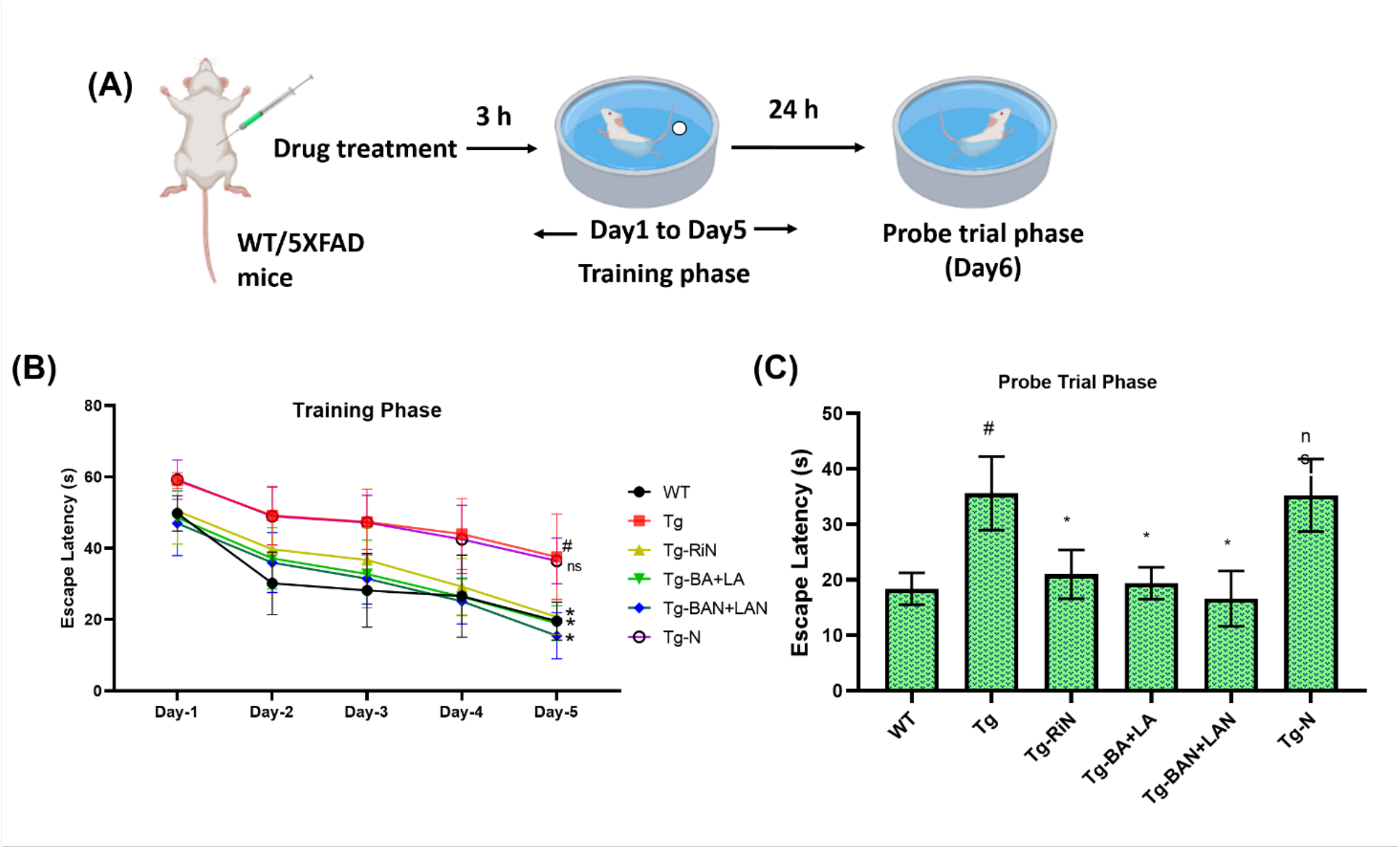
Amelioration of spatial memory impairment by nano-herbal formulation of bacosides and lauric acid efficaciously than nano-herbal formulation of rivastigmine in 5XFAD mice. (A) Experimental timeline of Morris water maze behavioral paradigm. (B) Analysis of 5 day training phase in different experimental groups. (C) Probe Trial analysis in different experimental groups. The treatment groups include wild type (WT) and 5XFAD mice. G*roup-1* Wild type treated with saline (WT), *group-2*5XFAD mice treated with Saline (Tg), *group-3*Tg treated with nano-formulation of rivastigmine (Tg-RiN), *group-4* bacosides and lauric acid (Tg-BA+LA), *group-5*Tg treated with nano-herbal formulation of BA and LA (Tg-BAN+LAN) and *group-6* Tg treated with nanoparticle alone (Tg-N). Histogram represents mean of the data (±SD). Statistical analysis were performed using One way Anova followed by Brown-Forsythe test (p< 0.05) between the groups (# - Tg vs WT, * - other treatment group vs Tg).

## Discussion

In the present study, we have delineated the therapeutic potential of a novel nano-herbal formulation of bacosides-lauric acid, BAN-LAN in preclinical AD transgenic 5XFAD mouse model. Combinatorial herbal formulation approach, nano-polymer encapsulation for better delivery, multi-targeted action of the bioactive and most importantly improved efficacy than conventional drug rivastigmine, all together in a single study confers diligence and novelty to our findings. Considering the multifactorial etiology and hypothesis of AD, we have evaluated recovery potential of BAN-LAN at cellular, molecular and behavioral level. We included AD candidate gene expression profiling, amyloid burden, neural atrophy, inflammation and spatial memory deficits, characteristic of neurodegeneration as experimental parameters in our study.

Previously, we reported that BAN-LAN combination attenuated scopolamine induced neurodegeneration in primary cortical neurons with concomitant upregulation of neurotrophins and memory markers of BDNF and Arc. In order to understand the effect of BAN-LAN as well as screen other control drug formulations in the context of AD, we employed Aβ-42 administered primary hippocampal neurons and investigated expression of AD candidate genes. Of note, we selected hippocampus in both our *in vitro* and *in vivo* experiments as hippocampal neurodegeneration precedes behavioral deficits in AD and is also responsible for short term memory loss, hall mark of AD (Vyas et al., 2020). Based on collection of published GWAS studies in AD (Andrews et al., 2020), we selected candidate genes of APP, PS1, ApoE and MAPT and others including apoptosis marker BAK, innate immune genes CD33 and CR1 that have been identified as novel risk genes for late onset AD; ADAM10, metalloproteinase acting in the non-amyloidogenic processing of APP and implicated in sporadic AD; ABCA7, crucial genetic determinant for late onset AD by regulating cholesterol metabolism and amyloid processing and clearance (Dib et al., 2021); CASS4, SNP’s of which showed significant correlation with pathological features of AD.

BAN-LAN attenuated Aβ42 induced changes in gene expression efficaciously than BA-LA as well as RiN for all the category of AD genes indicating its multi-targeted action. Also, the promising findings from primary neuronal culture data intrigued us to analyze the effects of BAN-LAN in AD transgenic mice *in vivo* and assess cellular and cognitive parameters, characteristic of AD related neurodegeneration. It is well established that Aβ induced senile plaque accumulation is a cardinal feature of AD and failure of its clearance leads to progression of the disease. Hence, drug discovery for AD has continuously aimed for inhibition of plaque formation as well as promotion of Aβ clearance. As Aβ peptide depositions are primarily observed hippocampus of AD patients (Zhang et al., 2011), we evaluated the effect of BAN-LAN and other drug formulations on hippocampal amyloid burden in AD mice. BAN-LAN significantly decreased Congo red-stained senile plaque numbers in 5XFAD mice and as anticipated was more potent than unconjugated BA-LA and nano-rivastigmine, RiN. Aβ is generated from APP through sequential cleavage by β-and γ-secretase (O’Brien and Wong, 2011) and our in vitro data showed that BAN-LAN efficaciously reduced the expression of Aβ induced APP mRNA than BA-LA and RiN indicating that it could be a possible molecular pathway.

Aβ accumulation in the brain of AD patients has been implicated in neuronal loss, synaptic dysfunction and memory impairment particularly in hippocampal regions (Bayer and Wirths, 2010).

5XFAD mice showed drastic hippocampal neurodegeneration as evident from loss of Nissl stained neurons though with regional variability; CA1 and CA3 showed greater extent of neuronal death than DG. Selective neuronal degeneration is a well-established feature of AD in which CA1 pyramidal cells degenerate in early onset AD whereas degeneration in other brain regions are observed in later stages (Saxena and Caroni, 2011; Fukutani et al., 2000). Interestingly, BAN-LAN treatment rescued neuronal death in all the hippocampal sub-regions indicating its potency to recover early as well as late onset AD associated neuronal atrophy and death. BAN-LAN showed efficiency in restoring neurodegeneration than BA-LA and RiN and even increased total number of neurons as compared to wild type indicating its capability to promote neurogenesis.

Increasing evidences suggest that AD pathogenesis is not limited to amyloid cascade hypothesis and neuronal death, rather it has strong association with the immunological processes in the brain (Heneka et al., 2014). In addition, innate immune genes being considered as novel AD risk genes and elevated level of inflammatory markers in AD human brain samples indicates that neuro-inflammation has crucial role in AD disease progression and severity (Leng et al., 2021). In light of these information, we investigated whether our drug formulations mainly BAN-LAN can reduce inflammation in the hippocampus of 5XFAD mice. Recent studies have suggested that reactive astrogliosis is pivotal for neuroinflammation and pathophysiology of AD. GFAP positive activated astrocytes can increase Aβ production and secretion of pro-inflammatory cytokines including TNF-alpha, IL-1B and IL-6 leading (Sajja et al., 2016) thereby leading to neuronal damage and synaptic dysfunction. BAN-LAN significantly reduced GFAP expression and thus astrocytic activation throughout the hippocampus of 5XFAD mice. However, GFAP expression was also reduced than wild type upon BAN-LAN treatment through reason still remains unexplored. It is important to mention here that although GFAP is a well-established marker of reactive astrogliosis, according to recent studies, it is only indicative of initial astrocyte reactivity (Kumar A et al., 2021) and other markers are needed to be examined to elucidate the complexity of astrocyte biology in AD pathogenesis.

Microglial proliferation and activation around amyloid plaques and associated neuroinflammation has also gained utmost prominence in AD pathophysiology (Hansen et al., 2018). Emerging evidences suggest that microglia have dual roles being involved in Aβ clearance by phagocytosis as well as its over-activation leading to inflammation and augmenting cognitive decline (Edler et al., 2021; Gomes-Leal, 2012). Abnormal microglial activation is also considered causal for AD development rather than a consequence (Hopperton et al., 2018). Here, we used elevated level of intracellular calcium binding protein, IBA-1 as the marker of microglia activation also reported in other studies. BAN-LAN substantially relieved microglia over-activation in hippocampus of 5XFAD mice and to the greatest extent in DG followed by CA3 and CA1. Spangenberg et al (2016) reported that elimination of microglia diminished neuroinflammation in 5XFAD mice, and is beneficial for hippocampal-dependent cognitive tasks. BAN-LAN also ameliorated hippocampal dependent spatial memory decline better than BA-LA and RiN in 5XFAD mice but more importantly it increased memory abilities even more than wild type indicating its role as a potential memory booster.

Multi-target drug action has often been ascribed to their epigenetic modifying property that act on transcriptional, post transcriptional and translational levels. Earlier, we reported that primary neurons challenged with amnesic drug scopolamine showed elevated levels of chromatin modifying enzymes, DNMT and HDAC while BAN-LAN significantly reversed the changes. Recently, Nativio et al (2020) reported that an integrated multi-omics approach identified crucial roles of epigenetic machinery for AD and pointed towards potential of epigenetic strategies for early detection and treatment. Therefore, exploring epigenetic targets of BAN-LAN in AD brain is essential to dissect the mechanism of action.

In summary, we showed that BAN-LAN novel nano-herbal formulation has wide spectrum of neuro-modulatory, anti-inflammatory and pro-cognitive properties ascribing to its preclinical efficacy in AD mouse model. Our formulation well qualifies with the current demand for multicomponent drugs with poly-pharmacological properties, minimal side effects and maximal availability in recovery of complex disorders like AD. Moreover, improved efficacy than rivastigmine paves path for its exploration in human iPSc models and later application in clinical trials. As mechanism of action has been a primary hurdle for clinical success of herbal compounds, BAN-LAN downstream molecular pathways in alleviating AD pathologies warrants further investigations.

Besides demonstrating the potency of natural products over synthetic chemicals, we report for that BA and LA encapsulated in nanoparticles (BAN and LAN) can promote the crucial processes of neuronal development. It is noteworthy that nanoencapsulation by lactoferrin conjugated polymersome can efficiently deliver high molecular weight herbal constituents to brain and thus are potent than their conventional counterparts. This can be attributed to the smaller size and low charge density makes the nanoparticles suitable for the evasion of endosomal/lysosomal pathway, hence increasing the cellular uptake through the lipid bi-layer via endocytosis.

## Acknowledgements

This study was supported by a grant and fellowship of Dr. Ashish Kumar from Cognitive Science Research Initiative Scheme, Department of Science & Technology, Govt of India (SR/CSRI-P1/2017/41). The authors acknowledge Dr Beena Pillai, CSIR-IGIB, India for extending laboratory facilities required for histology and imaging related experiments. The authors also thank the Nanoscale Research Facility, IIT Delhi for their advanced instrumentation facility.

